# Taxonomic and biogeographic insights into Australasian fingerworts in the *Lepidozia ulothrix* clade

**DOI:** 10.64898/2026.09.09.750534

**Authors:** Antonio L. Rayos, Simon Y. W. Ho, Matthew A. M. Renner

## Abstract

The floras of Australia and New Zealand are characterised by very high endemism of bryophytes. The highly diverse genus *Lepidozia* (fingerworts; family Lepidoziaceae) is represented by more than 20 mostly endemic species in these countries. The genus has been monographed in New Zealand, but issues surrounding species circumscription remain. This study focused on the *Lepidozia ulothrix* clade, with members shared between Australia and New Zealand. We performed phylogenetic analyses of a multilocus data set and applied several methods for molecular species delimitation. Our results support the conspecificity of *L*. *serrulata* and *L*. *ulothrix*, which is compatible with patterns of morphological variation observed in Tasmanian and New Zealand populations. The distribution of *L*. *septemfida* extends to the Wet Tropics Bioregion of north-east Queensland, where the species is represented by a distinct subspecies rather than a completely different species. The morphological differences between the Queensland and south-east Australian populations, as well as the inconsistency among the species-delimitation results, support the division of the species into two subspecies. Tasmanian and Victorian specimens previously attributed to the New Zealand endemic *L*. *hirta* belong to a distinct and unnamed lineage. Our study shows that integrative taxonomic approaches can uncover overlooked diversity and refine existing species circumscriptions, even in relatively well-studied Australasian lineages such as *Lepidozia*.

## Introduction

Liverworts are found in a wide variety of habitats throughout the world, including the six floristic kingdoms that have been recognised for bryophytes (Schofield 1992). Endemism is notably high in the Holantarctic, which includes Australia, New Zealand, and temperate southern South America. New Zealand harbours 52% endemic species of liverworts (Vanderpoorten and Goffinet 2009), while similar rates of endemism characterise the floras of Australia (Stevenson et al. 2013) and southern South America (Gradstein et al. 2001). Several species-rich families of liverworts have centres of species and phylogenetic diversity in this region. These include the families Lophocoleaceae and Lepidoziaceae, both of which contribute to the rich ‘mossy’ character of southern temperate rainforest habitats.

The Lepidoziaceae are the third-largest liverwort family by number of genera, with between 27 and 30 currently accepted (Schuster 2000), almost all of which occur in the Holantarctic. The type genus of the family is *Lepidozia* (Dumort.) Dumort., commonly known as ‘fingerworts’ (a term shared with *Kurzia*; Edwards 2020). The species in this genus show diverse ecology, principally occurring on soil, peat, or decaying logs (Engel and Glenny 2008). Some species also grow on soil-covered rocks, while a few can invade lower trunks of trees. Many species reach the alpine zone where they occupy damp sites of alpine grasslands or soil in crevices of crags or cliffs.

In the most recent phylogenetic study of Lepidoziacaeae (Rayos et al. 2024), it was inferred that *Lepidozia* diverged from *Neolepidozia*, its sister genus, 74 (95% CI 94–55) million years ago (MYA). This is the most recent generic divergence in the family. *Lepidozia* comprises around 80 species worldwide and is among the largest genera in the family by number of species (Engel and Glenny 2008). Twenty-three species of *Lepidozia* occur in New Zealand, of which 16 are endemic, and 27 occur in Australia (Renner et al. 2024). However, difficulties in species delimitation have led to continued uncertainty in the number of species in both countries.

New Zealand *Lepidozia* was subject of a dedicated revision (Engel and Schuster 2001) and follow-up contribution (Engel 2004) in preparation for the Liverwort Flora of New Zealand project (Engel and Glenny 2008). However, subsequent molecular phylogenetic studies that included a haphazard sampling of multiple representatives of species found that some, including *L. pendulina* (Hook.) Lindenb. and *L. microphylla* Hook., were non-monophyletic. Further investigations resulted in the formal recognition of *L. bragginsiana* (Cooper and Renner 2014) and *L. nanophylla* (Renner 2025). However, other species circumscription issues raised by the study of Cooper et al. (2011, 2012), associated with *Lepidozia obtusiloba* and *L. ulothrix*, have either not yet been addressed or have been only partially resolved.

*Lepidozia ulothrix*, as circumscribed by Engel and Glenny (2008), included plants from mainland Australia, Tasmania, and New Zealand. However, individuals from mainland south-east Australia formed a distinct sister group to the remainder of the *L. ulothrix* clade, rather than being resolved as a sister group to the lineage containing individuals from Tasmania and New Zealand (Cooper et al. 2012). The lineage from Tasmania and New Zealand corresponded to *L. ulothrix* and *L. serrulata,* while the lineage from mainland Australia corresponded to *L. septemfida* (Cooper et al. 2012, 2013). *Lepidozia septemfida* is morphologically distinct from *L. ulothrix* in the large laciniate teeth on the dorsal leaf margin. *Lepidozia septemfida* was formally reinstated to reflect the morphological and phylogenetic distinctiveness of plants from mainland Australia by Cooper et al. (2013), who noted that all Australian mainland specimens of the *L. ulothrix* clade examined as part of their study belonged to *L. septemfida*. However, the distribution of *L. septemfida* on mainland Australia remains incompletely documented.

When *L. serrulata* was proposed as a new species, a close relationship between *L. serrulata* and *L. hirta* was inferred by Engel (2004) on the basis of their similar leaf armature and other distinct morphological features. *Lepidozia serrulata* was placed in sect. *Kirkii* with *L. kirkii* and *L. hirta*, rather than in sect. *Austrolepidozia* with *L. ulothrix*(Engel 2004). However, two specimens determined as *L. serrulata* were resolved among *L. ulothrix* in the phylogeny of Cooper et al. (2012), suggesting that *L. serrulata* is, in fact, closer to *L. ulothrix*, consistent with their shared possession of caudate leaf lobe apices comprising elongate cells, among other characters that distinguish them from *L. hirta* and *L. kirkii* (Engel and Schuster 2001; Engel 2004). The close relationship between *L. serrulata* and *L. ulothrix* (Cooper et al. 2012) suggests a need for further investigation into the distribution and circumscription of these two species. *Lepidozia serrulata* was not listed as part of the Australian flora in the recently updated checklist of Australian liverworts and hornworts because the species was regarded as a New Zealand endemic by Engel and Glenny (2008), and more investigation of Tasmanian plants attributed to *L. serrulata* was required (Renner et al. 2024).

In this study, we use molecular phylogenetic analysis and other tools to provide a framework for clarifying the taxonomy and distribution of the Australasian fingerworts of the *L*. *ulothrix* clade. We include *L. hirta*, another species in the *L. ulothrix* clade that was reported for Australia by Meagher (2006) but was listed as a New Zealand endemic (Engel and Glenny 2008). The results of our study demonstrate how integrating molecular data with morphological observations can benefit in resolving taxonomic issues in *Lepidozia* and other challenging taxa.

## Materials and Methods

### Molecular data set

We sampled 13 herbarium specimens of *Lepidozia* from four herbaria in Australasia (Supplementary Table 1). From each sample, about 25 mg of dried tissue was cleaned and sent to the Australian Genome Research Facility (Brisbane) for DNA extraction, PCR amplification, and amplicon purification. Dual-direction Sanger sequencing of four markers (chloroplast *rbcL* and *trnL*–*trnF*, mitochondrial *nad5*–*nad4*, and nuclear *ITS2*) by DNA BDT labelling reaction and capillary separation was carried out on an Applied Biosystems 3730xl Genetic Analyzer (Supplementary Table 2). We excluded sequences that did not show close affinity with available sequences from *Lepidozia*, as assessed using BLASTn searches. We combined the resulting 43 sequences with available sequence data from 78 individuals representing 41 species of *Lepidozia*, plus 27 individuals representing 11 species of *Neolepidozia* as the outgroup, to produce a data set comprising a total of 118 taxa. The outgroup *Neolepidozia* is the sister genus to *Lepidozia* (Cooper et al. 2012; Rayos et al. 2024).

We aligned the sequences of the four markers individually using MAFFT (Katoh and Standley 2013) and removed poorly aligned regions using trimAl with the automated algorithm optimised for maximum-likelihood analyses (Capella-Gutiérrez et al. 2009). Our assembled data supermatrix had an occupancy of 74.79%, with 353 sequences available out of a possible 472 (four markers from 118 individuals; Supplementary Table 3).

For 94 samples, including all of those in this study, we attempted to generate a genome-wide panel of single-nucleotide polymorphisms to enable estimation of population structure and gene flow. However, we found that most of the obtained sequence data were from contaminants including bacteria and viruses, and we failed to obtain sufficient useable data to allow reliable analyses. This can partly be attributed to degraded DNA from the herbarium samples.

### Phylogenetic analyses

We performed a phylogenetic analysis of the concatenated data set using maximum likelihood in IQ-TREE 2 (Bui et al. 2020), with the best-fitting partitioning scheme selected using a greedy search. Data subsets were allowed to evolve at different relative rates, such that the branch lengths were proportionate across subsets (Duchêne et al. 2020). Node support values were estimated using 1000 bootstrap replicates. We also estimated the phylogeny using Bayesian phylogenetic analysis in BEAST v2.7.3 (Bouckaert et al. 2019) using a birth-death tree prior and uncorrelated lognormal relaxed clock (Drummond et al. 2006), allowing a separate relative substitution rate for each data subset (partitioned according to the scheme selected in IQ-TREE). The posterior distribution was estimated using Markov chain Monte Carlo sampling, with samples logged every 5000 steps over a total of 50 million steps. We then performed the same analysis using only the two chloroplast DNA markers, without partitioning, to infer a tree for our subsequent species-delimitation analyses.

In another analysis, we inferred separate trees for the chloroplast, mitochondrial, and nuclear markers using IQ-TREE 2 with substitution model selection. Using ASTRAL III (Zhang et al. 2018), we inferred the species tree from these three gene trees. We omitted two samples that caused low support values in the clade containing them because these samples lacked sequence data for two out of the four markers. In the final data set of 116 samples, the Australasian *Lepidozia ulothrix* species group was represented by 23 individuals sampled throughout their known geographic range (Figure 1).

**Figure 1.**
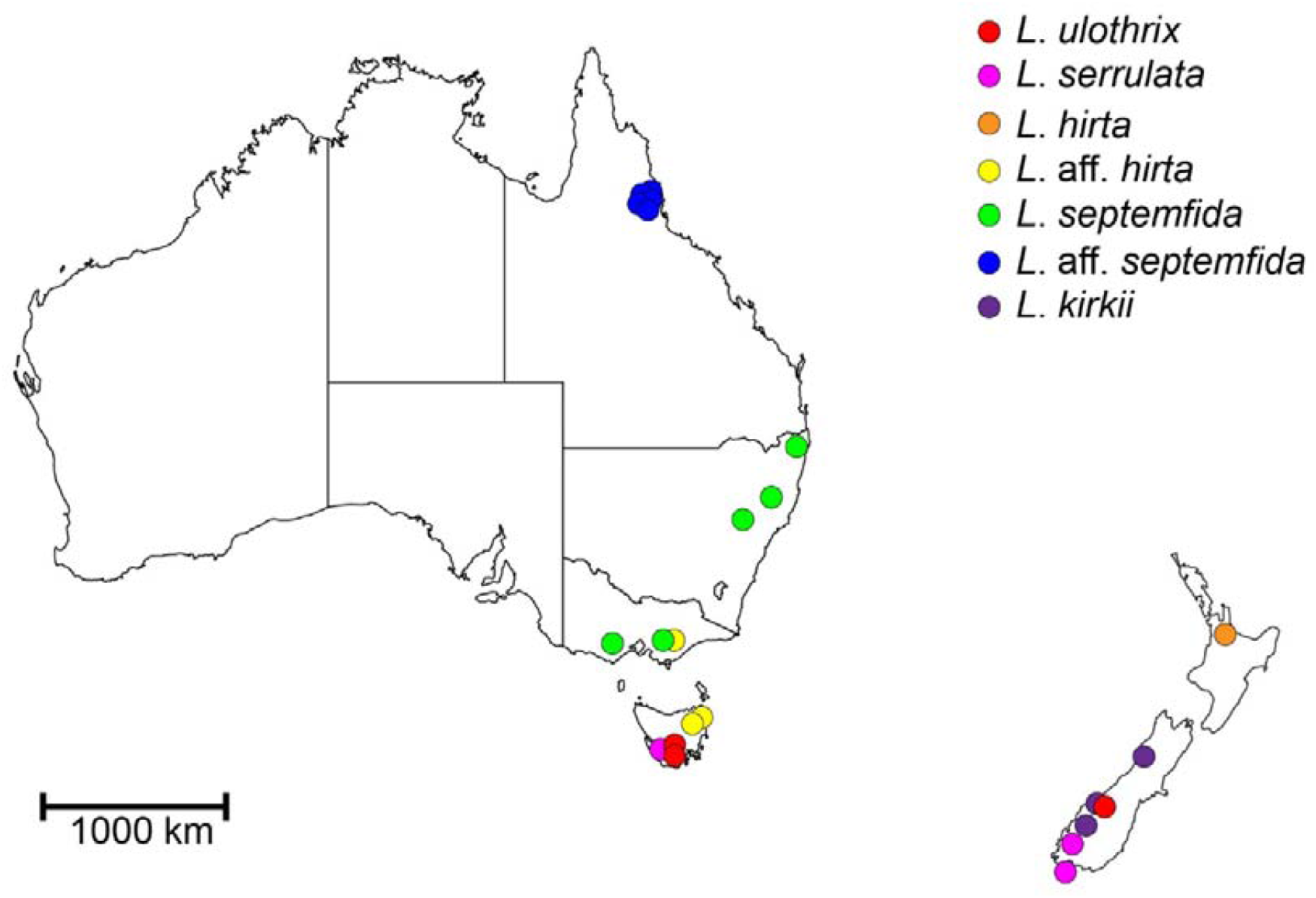
Map of Australia and New Zealand showing the provenance of the 23 samples representing the *L*. *ulothrix* species group in this study.

### Species-delimitation analyses

We employed three methods for molecular species delimitation: the Generalized Mixed Yule Coalescent (GMYC) approach (Fujisawa and Barraclough 2013), the Bayesian implementation of the Poisson tree processes (bPTP) model (Zhang et al. 2013), and Automatic Barcode Gap Detection (ABGD; Puillandre et al. 2012). The first two methods use phylogenetic trees to propose species delimitations. In our GMYC analysis, we used the single-threshold (S-GMYC) and multiple-threshold (M-GMYC) methods and analysed the Bayesian tree inferred from the two chloroplast markers (with the outgroup removed). In bPTP maximum likelihood (bPTP-ML), we used the chloroplast tree (with the outgroup removed) inferred using IQ-TREE and performed the analysis with the default settings. We ran each of these methods using the respective web server.

In our molecular species delimitation using ABGD, we performed the analysis using default settings except for the value of X (proxy for minimum gap width) set to 0.9 in ABGDpy 0.1 (Vences et al. 2021). We used an edited sequence alignment (comprising the four concatenated markers), from which we removed the outgroup taxa and some ingroup taxa to ensure that all pairwise comparisons involved overlapping sequence data.

## Results

### Phylogeny

Our phylogenetic analyses of the chloroplast and nuclear data yielded trees that are mostly congruent. However, only the chloroplast tree supports monophyly of the *L*. *ulothrix* species group. The mitochondrial tree, which has a high degree of incongruence with the two other trees, contains several polytomies; these can be attributed to a lack of informative sites in the chosen marker. The species tree inferred using a summary-coalescent approach, although having poor support for most nodes, is substantially congruent with the well-resolved maximum-likelihood and Bayesian trees (Figure 2A).

**Figure 2.**
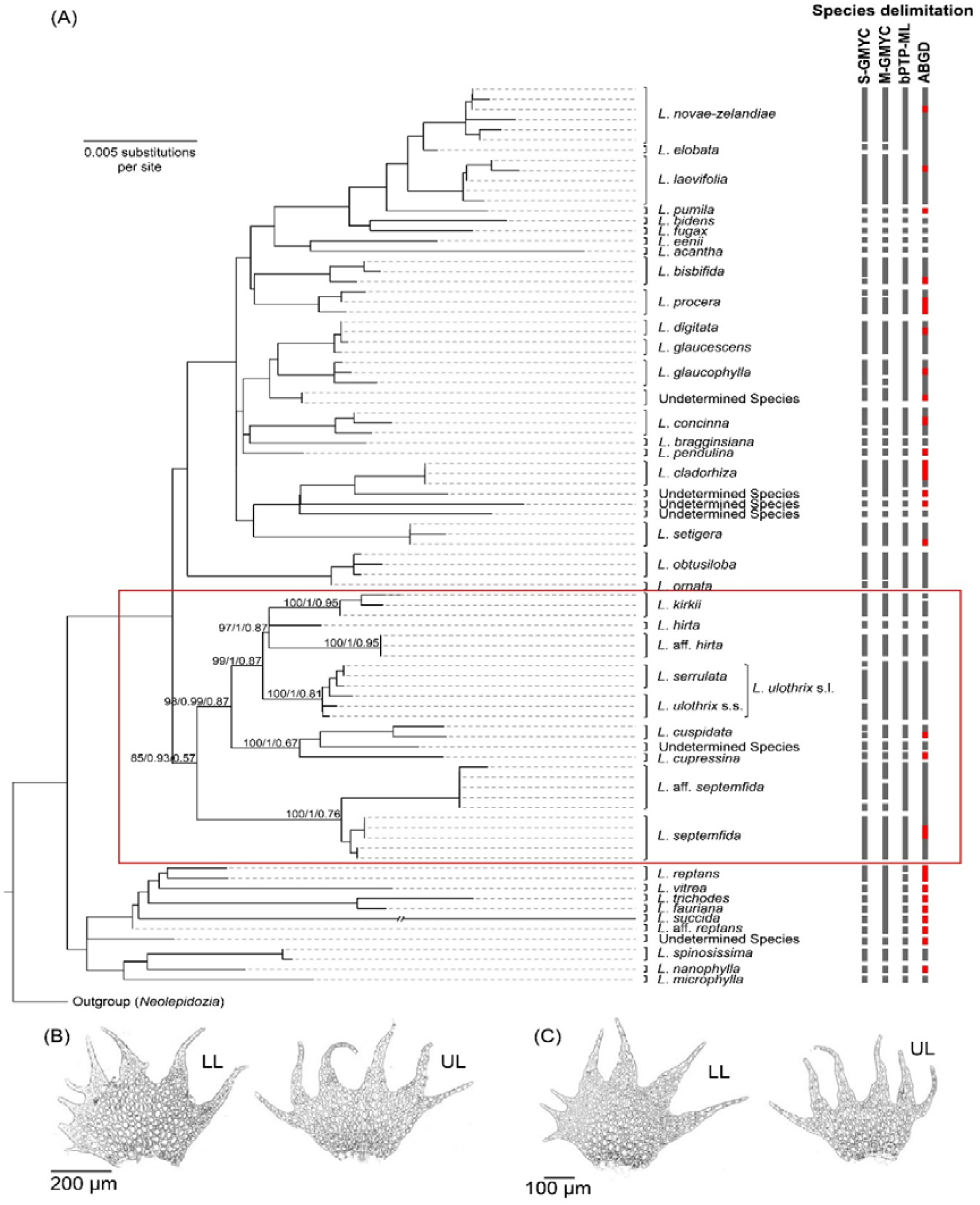
(A) Maximum likelihood tree of *Lepidozia* inferred using four markers. The portion inside the red box includes the *L*. *ulothrix* species group, shown with node support values from maximum-likelihood, Bayesian, and summary-coalescent phylogenetic analyses. The terminal branch (0.0318 substitutions per site) leading to *L*. *succida* has been truncated. The outgroup, comprising 27 samples from the genus *Neolepidozia*, has been pruned from the tree to improve visualisation of the ingroup. Vertical grey bars show the species-delimitation results from GMYC (single and multiple threshold), bPTP-ML, and ABGD methods. Vertical red bars indicate the samples removed from the analysis to ensure overlap of sequences among all samples. (B) Lateral leaves (LL) and underleaves (UL) of the undescribed species resembling *L*. *hirta*. (C) Lateral leaves (LL) and underleaves (UL) of the new subspecies of *L*. *septemfida* in Queensland.

The six samples representing *L*. *ulothrix* and *L*. *serrulata* appeared in a clade with relatively short terminal branches. The three specimens identified as *L*. *hirta* from Victoria and Tasmania formed a distinct clade separate from the samples of *L*. *hirta* from New Zealand. The nodes associated with the abovementioned clades have bootstrap support values of 100% and posterior probabilities of 1.00. The five specimens from Queensland that are morphologically similar to *L*. *septemfida* formed a sister clade to this species. However, the support values for the node associated with this clade are relatively low, while the clade containing these Queensland specimens and all of the specimens of *L*. *septemfida* is well-supported.

### Species delimitation

The three species-delimitation methods GMYC, bPTP-ML, and ABGD yielded consistent results in many cases (Figure 2A). *Lepidozia setigera*, represented by three samples, was consistently delimited by all three methods. In contrast, the samples representing *L*. *digitata* and *L*. *glaucescens* were consistently inferred not to be distinct species. The molecular species delimitation yielded conflicting results in some cases. For instance, *L*. *procera* was delimited as a distinct species by bPTP-ML and ABGD but not by GMYC. The single– and multiple-threshold methods of GMYC yielded 49 and 41 putative species, respectively, while bPTP-ML yielded only 39. None of these is perfectly consistent with the currently defined species.

Within the *L*. *ulothrix* species group, the three samples that superficially look like *L*. *hirta* were consistently delimited as a distinct species. This was also the case with the six samples representing *L*. *ulothrix* and *L*. *serrulata* (except when we used the S-GMYC method, which tends to oversplit species), thus implying that they are conspecific. In contrast, the three samples of *L*. *kirkii* were delimited as a species only by GMYC and bPTP-ML. The Queensland samples having strong morphological affinity to *L*. *septemfida* were delimited as a distinct species only by bPTP-ML, whereas *L*. *septemfida* was delimited only by GMYC and bPTP-ML. The inconsistent delimitation of *L. septemfida* across the three methods suggests that the Queensland samples could be a distinct subspecies of *L*. *septemfida* rather than a separate species.

## Discussion

The results of our phylogenetic analyses and molecular species delimitation have provided various insights into the taxonomy and biogeography of the Australasian members of the *L*. *ulothrix* species group. We are able to deduce species boundaries based on these results, although interpretation depends on the choice of species concept. The biological species concept, which defines species as groups of populations that can interbreed and produce fertile offspring (Mayr 1942), is the most widely accepted but is difficult to apply in this study because it is mainly based on tokogenic hypotheses rather than phylogenetic evidence (Fitzhugh 2005). The morphological species concept, which defines species as the smallest group of organisms that are consistently distinct and easily distinguishable from other groups (Cronquist 1978), is also widely accepted and can be applied to the species in this study, but it can also be prone to biases. Therefore, using it in conjunction with another species concept is necessary. Molecular species delimitation employs the phylogenetic species concept, which defines species as the smallest diagnosable cluster of individuals in an inferred evolutionary tree (Cracraft 1983). We applied this species concept in interpreting the results of the phylogenetic analyses and molecular species delimitation. We also applied the morphological species concept to further support our interpretations.

### *Conspecificity of* L. ulothrix *and* L. serrulata

Our results suggest that *L*. *ulothrix* and *L*. *serrulata* are different morphological expressions of the same variable species, which occurs in New Zealand and Tasmania. This is supported in the maximum-likelihood tree by the relatively short terminal branches among the tips representing those two species, which are comparable to the branches within the clades containing individuals of the same species (e.g., *L*. *novae-zelandiae* and *L. laevifolia*). Moreover, the six samples representing these species were consistently grouped into a single species by the M-GMYC, bPTP-ML, and ABGD methods, which supports the conspecificity of the two taxa. Features distinguishing *L*. *serrulata* from *L*. *ulothrix* include pale yellow-green leaves, bipinnately branched shoot systems, entire leaf lobes, and denticulate-dentate leaf disc margins. Both *L. serrulata* and *L. ulothrix* share ciliform lobe apices comprising elongated cells on both leaves and underleaves. Following the principle of priority, the name *L*. *ulothrix* is the one to be retained as the accepted name, while *L*. *serrulata* is the one to become a synonym.

*Lepidozia serrulata* occurs as ground-dweller in protected, non-exposed sites and in mosaic communities of stagnant ponds (Engel and Glenny 2008). In contrast, *L*. *ulothrix* s.s. occurs in more open habitats. In leafy liverworts, slight variations in branching can provide a basis for adaptation during the colonisation of new environments (Schuster 1984). This can at least partially explain the difference in branching between these two entities because they occupy contrasting habitats. There has never been any published study of how environment can affect the branching pattern in leafy liverworts. It is possible that *L*. *serrulata* or *L*. *ulothrix* s.s. can exhibit phenoplasticity for this trait depending on the habitat. However, this requires a thorough investigation.

### New species from South-east Australia

Three collections from Tasmania and Victoria determined as *L*. *hirta* represent an undescribed species related to *L*. *hirta* and *L. kirkii*. This result is strongly supported by our phylogenetic analyses and the three species-delimitation methods, confirming that *L*. *hirta* is a New Zealand endemic. The new species slightly differs from *L*. *hirta* in morphology of first branch leaves and underleaves (Figure 2B). The former has less dissected lobes in both leaves and underleaves than the latter. More details will be presented in the species description in a separate article.

### New subspecific taxon under L. septemfida

*Lepidozia septemfida*, when first published (Stephani 1909), was known to occur only in New South Wales and was noted to be morphologically similar to *L*. *ulothrix*. However, previous phylogenetic studies (Cooper et al. 2012; Rayos et al. 2024) and this study showed that *L*. *septemfida* is more closely related to extra-Australasian species than to *L*. *ulothrix*. The samples of *L*. *septemfida* from New South Wales and Victoria formed distinct geographic clades with high support values. The samples resembling this species collected from Queensland (Figure 2C) also formed a distinct clade. To decide whether or not these represent an undescribed species, we considered the results from the species-delimitation methods. The samples of *L*. *septemfida* from New South Wales and Victoria were delimited as a distinct species by GMYC and bPTP-ML, but not by ABGD which grouped them together with the samples from Queensland. The latter samples were delimited as a separate species only by bPTP-ML. The inconsistency in these molecular species delimitations raises the possibility that the Queensland population represents a distinct subspecies of *L*. *septemfida*. The populations in New South Wales and Victoria likely represent another subspecies, as implied by their geographic distinctness. The morphological differences between the two putative subspecies include the difference in the size of the teeth in the dorsal margin of the first branch lateral leaves. Details on morphology will be provided in a separate article where these subspecies will be proposed.

An alternative explanation for the inconsistency in the molecular species delimitations for these samples is that the two species are still at an early stage of divergence. Morphological diagnosability and reciprocal monophyly are among the properties that provide evidence of lineage separation, but these properties are not necessarily useful for species delimitation in the early stages of divergence (de Queiroz 2007). The ABGD method, which placed all of the samples of *L*. *septemfida* as a distinct species, is not suitable for inferring speciation events that occurred very recently (Puillandre et al. 2012). Both GMYC and bPTP-ML, which delimited the samples of *L*. *septemfida* from New South Wales and Victoria as a separate entity from the Queensland population, have been shown to be sensitive to the time since divergence (Luo et al. 2018). Nonetheless, the molecular data that we gathered do not provide sufficient evidence to show that the New South Wales–Victoria and Queensland populations are separate species. Although the reciprocal monophyly of the two populations hints at reproductive isolation, genome-wide data will be needed to quantify the level of gene flow between the two populations.

### Biogeography of the L. ulothrix species group

Within the *L*. *ulothrix* species group, there is a distinct clade of extra-Australasian species (from Africa and South America) that forms a sister lineage to the rest of the species group except *L*. *septemfida* s.l. In the most recent molecular dating study of Lepidoziaceae (Rayos et al. 2024), it was estimated that the crown age of *Lepidozia* is about 63 (95% CI 83–47) MY, which predates the final split of Australia from Antarctica about 45 MYA (van den Ende et al. 2017). However, the node associated with the *L*. *ulothrix* clade is about 33 (95% CI 46–22) MY, so the evolutionary relationship of the extra-Australasian members of the clade to the rest of the species group cannot be readily explained by plate tectonics. Further studies with denser species sampling, including more taxa from Africa and South America, will shed light on the deeper evolutionary history of the genus.

## Conclusions

Using a combination of molecular phylogenetic analyses and species-delimitation methods, we have been able to provide taxonomic and biogeographic insights into the Australasian members of the *L*. *ulothrix* species group of fingerworts. Our study shows how integrative approaches in taxonomy can aid in discoveries of undocumented diversity in taxonomically challenging taxa such as *Lepidozia*. More species groups of the genus in Australasia possibly need further investigation, along similar lines to those used in this study. Furthermore, our findings here can contribute valuable information for revising floras and checklists of *Lepidozia* species in Australasia.

## Acknowledgements

The authors are grateful to the following people for their kind assistance during the acquisition of the samples used in this study: Brendan Lepschi, Christine Cargill, and Judith Curnow of the Australian National Herbarium (Canberra); Hannah McPherson, Yola Metti, and Margaret Heslewood of the National Herbarium of New South Wales (Sydney); Nimal Karunajeewa, Niels Klazenga, and Rebecca Le Get of the National Herbarium of Victoria (Melbourne); and Dhahara Ranatunga, Dan Blanchon, and Yumiko Baba of the Auckland War Memorial Museum Herbarium (Auckland). A.L.R. was supported by postgraduate student travel grants administered by the Australasian Systematic Botany Society Inc. and Society of Australian Systematic Biologists (both supported by the Australian Biological Resources Study); by a Graduate Student Research Award from the Society of Systematic Biologists; and by the Foreign Graduate Scholarship Program from the Science Education Institute of the Department of Science and Technology (DOST-SEI) of the Philippines.

## Data Availability Statement

The DNA sequences that support the findings of this study are available in GenBank of NCBI at https://www.ncbi.nlm.nih.gov/ under the accession numbers provided in Supplementary Table 3.

---

**Supplementary Table 1.**
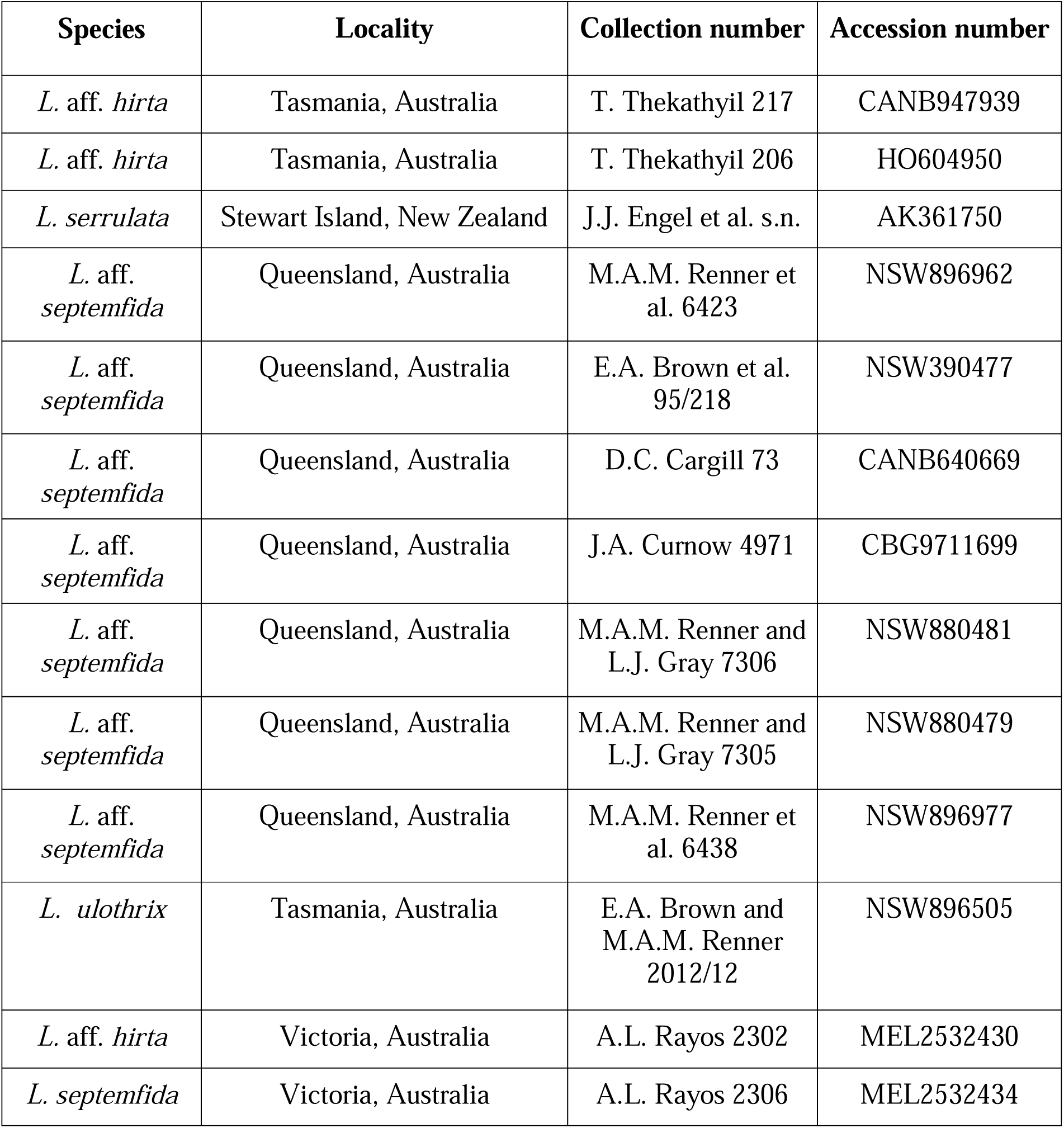
Details of herbarium samples of *Lepidozia* used in this study.

**Supplementary Table 2.**
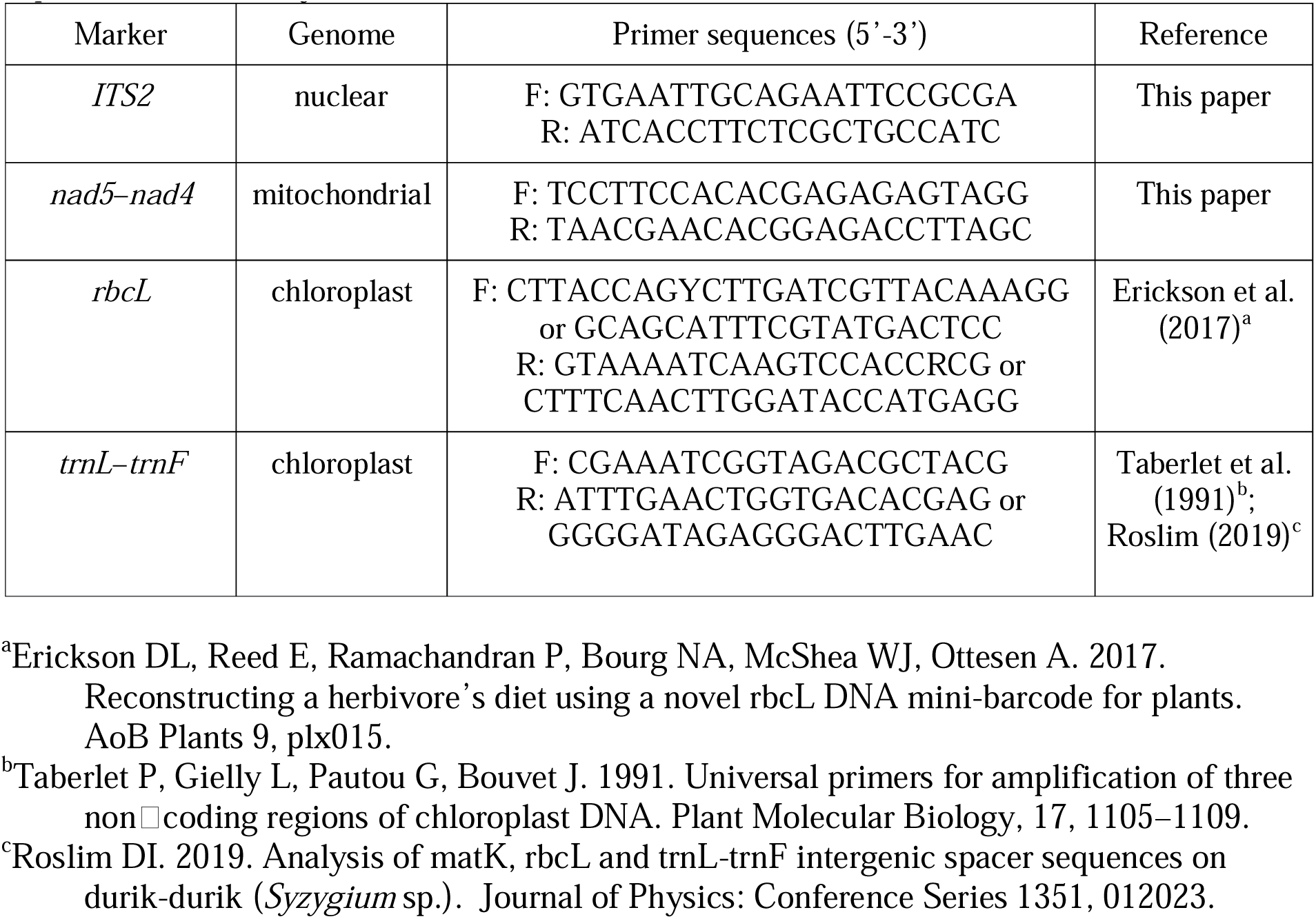
Molecular markers amplified and sequenced for the samples of *Lepidozia* in this study.

**Supplementary Table 3.**
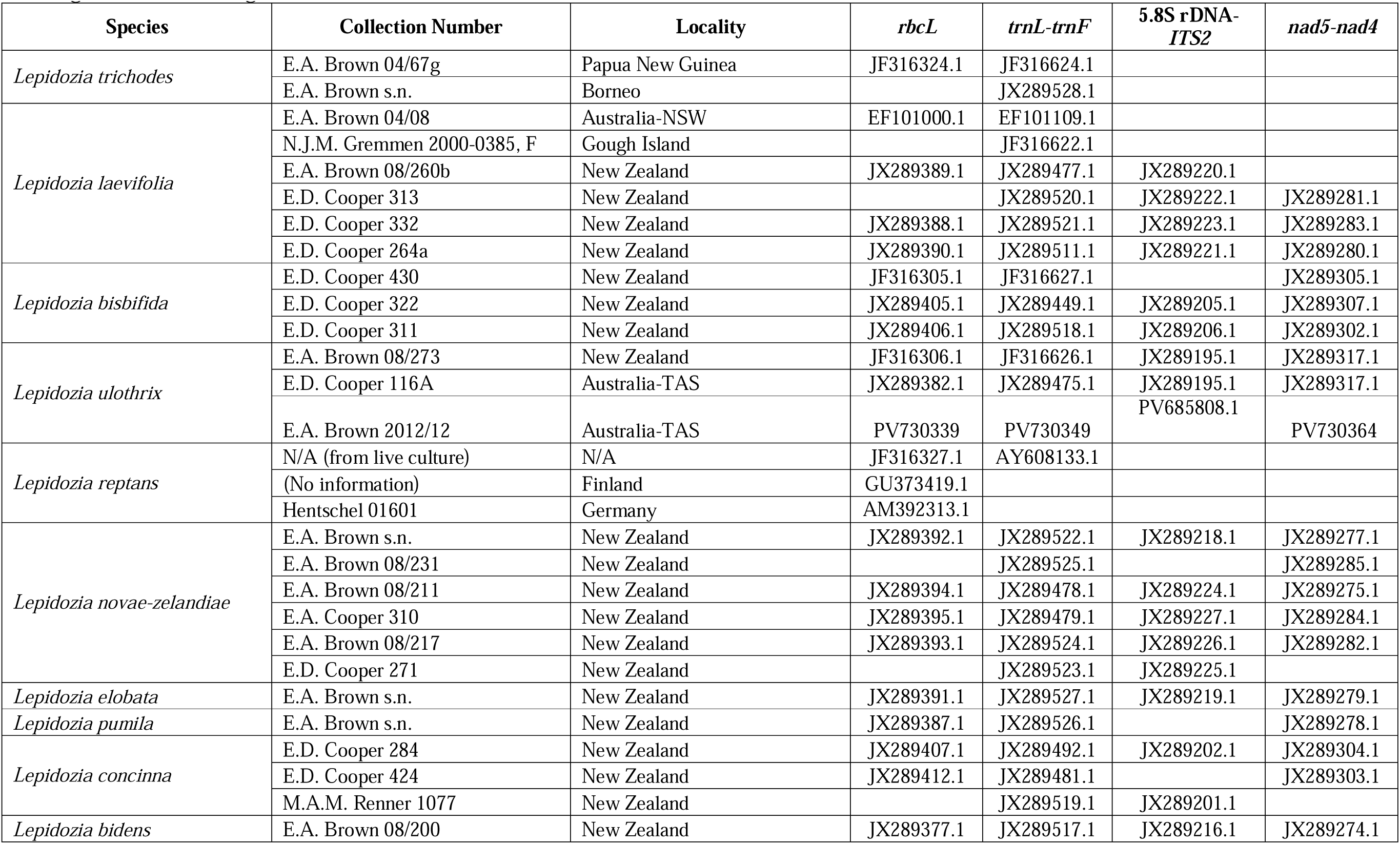
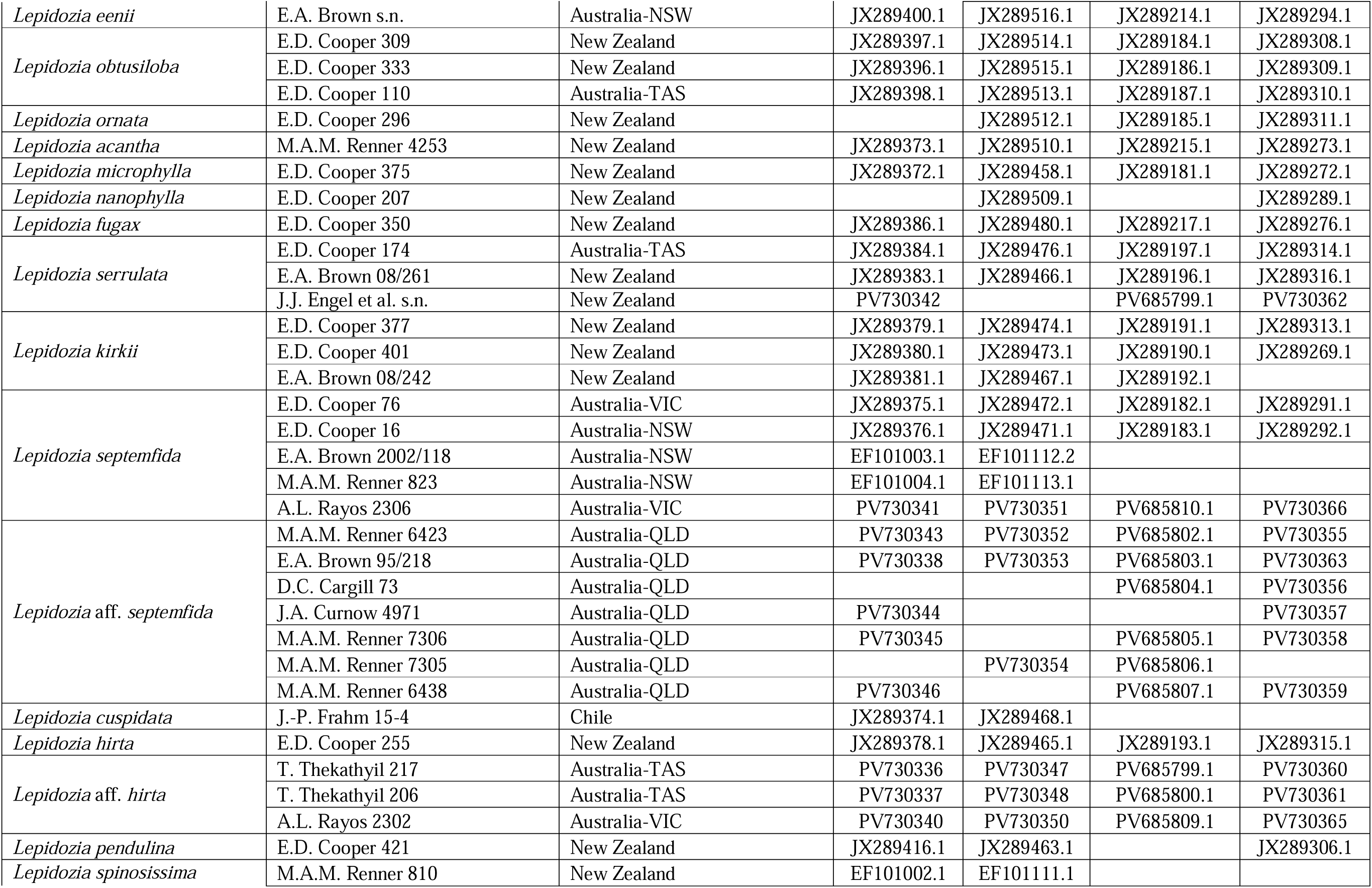
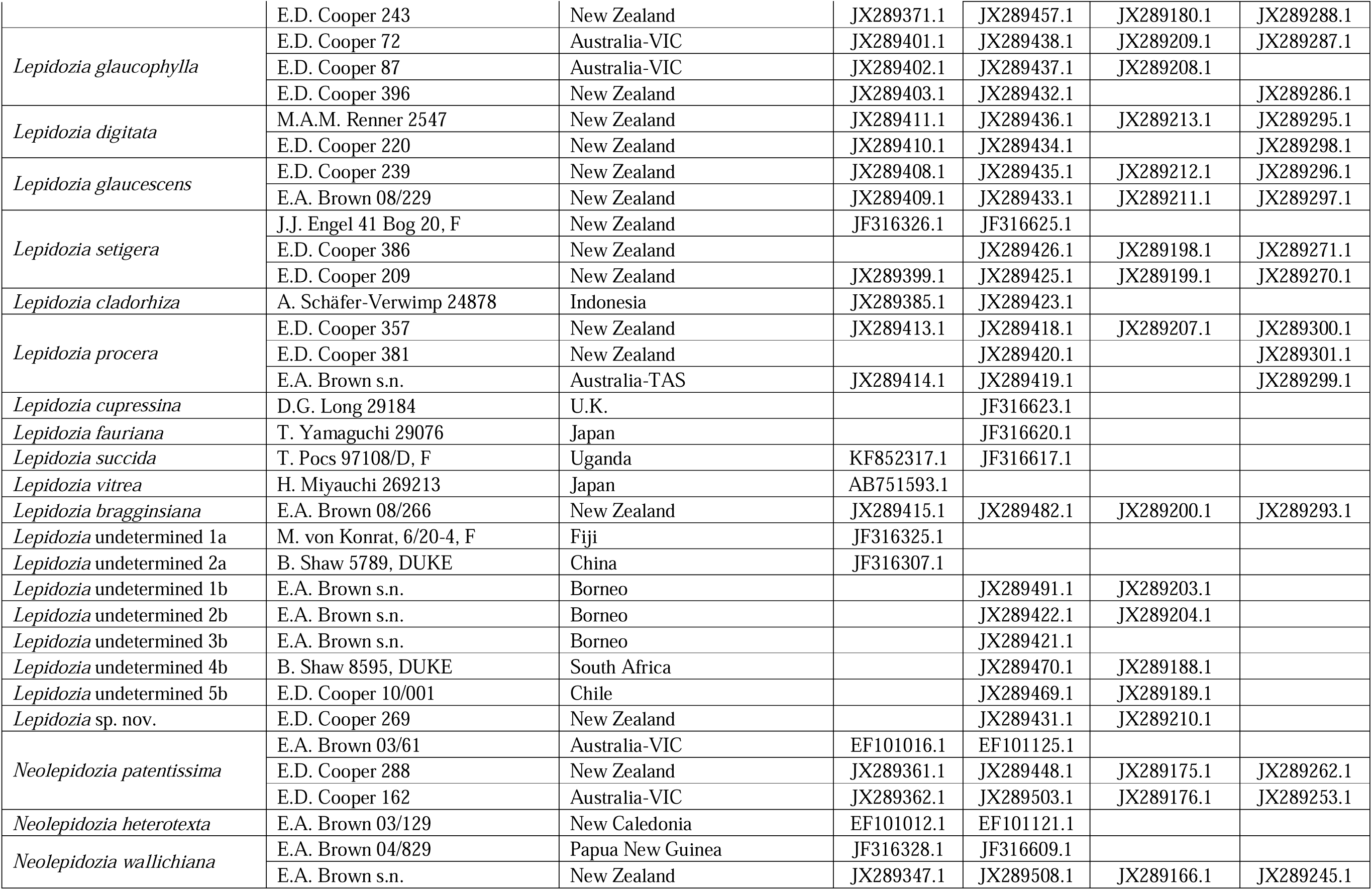
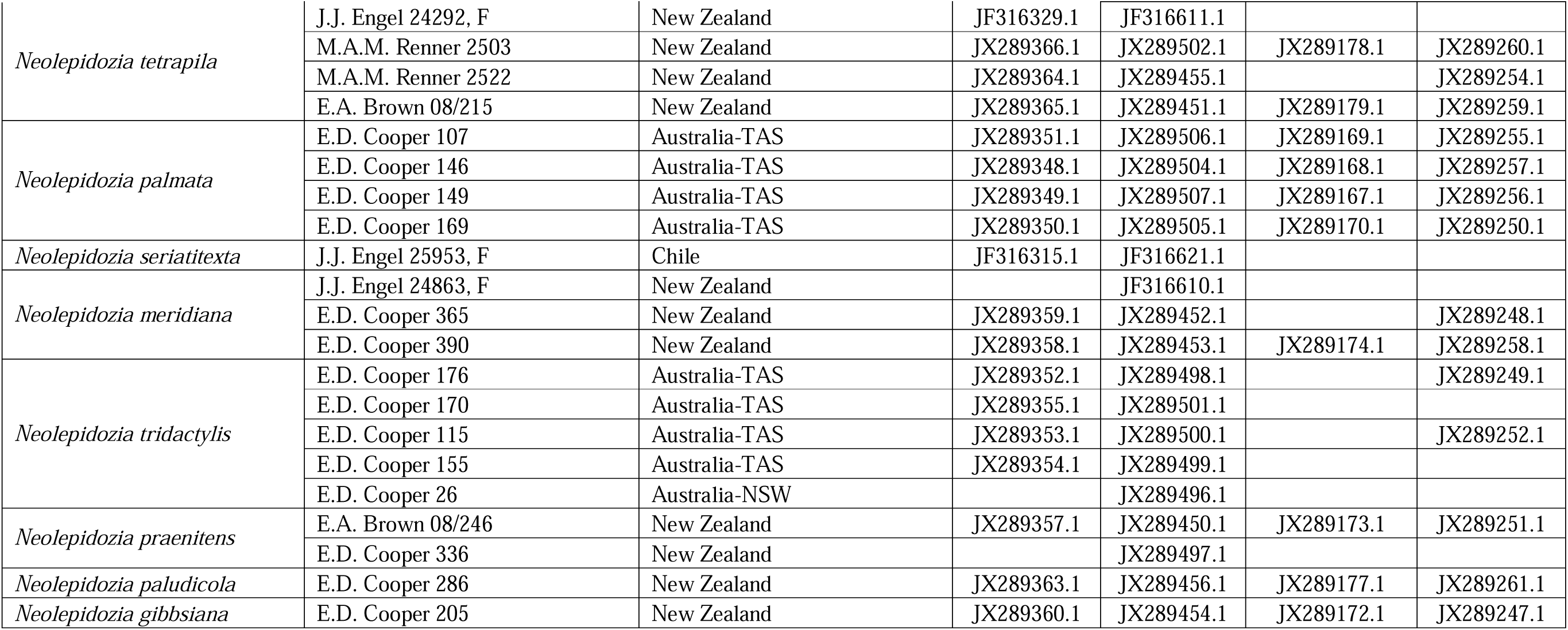
The 118 samples of *Lepidozia* and *Neolepidozia* (outgroup) analysed in this study, with GenBank accession numbers given for the four genetic markers.

