## Supplementary Tables for "Taxonomic and biogeographic insights into Australasian fingerworts in the *Lepidozia ulothrix* clade"

**Supplementary Table 1.** Details of herbarium samples of *Lepidozia* used in this study.

| **Species** | **Locality** | **Collection number** | **Accession number** |
| --- | --- | --- | --- |
| *L.* aff. *hirta* | Tasmania, Australia | T. Thekathyil 217 | CANB947939 |
| *L.* aff. *hirta* | Tasmania, Australia | T. Thekathyil 206 | HO604950 |
| *L. serrulata* | Stewart Island, New Zealand | J.J. Engel et al. s.n. | AK361750 |
| *L.* aff. *septemfida* | Queensland, Australia | M.A.M. Renner et al. 6423 | NSW896962 |
| *L.* aff. *septemfida* | Queensland, Australia | E.A. Brown et al. 95/218 | NSW390477 |
| *L.* aff. *septemfida* | Queensland, Australia | D.C. Cargill 73 | CANB640669 |
| *L.* aff. *septemfida* | Queensland, Australia | J.A. Curnow 4971 | CBG9711699 |
| *L.* aff. *septemfida* | Queensland, Australia | M.A.M. Renner and L.J. Gray 7306 | NSW880481 |
| *L.* aff. *septemfida* | Queensland, Australia | M.A.M. Renner and L.J. Gray 7305 | NSW880479 |
| *L.* aff. *septemfida* | Queensland, Australia | M.A.M. Renner et al. 6438 | NSW896977 |
| *L. ulothrix* | Tasmania, Australia | E.A. Brown and M.A.M. Renner 2012/12 | NSW896505 |
| *L.* aff. *hirta* | Victoria, Australia | A.L. Rayos 2302 | MEL2532430 |
| *L. septemfida* | Victoria, Australia | A.L. Rayos 2306 | MEL2532434 |

**Supplementary Table 2.** Molecular markers amplified and sequenced for the samples of *Lepidozia* in this study.

| Marker | Genome | Primer sequences (5’-3’) | Reference |
| --- | --- | --- | --- |
| *ITS2* | nuclear | F: GTGAATTGCAGAATTCCGCGA  R: ATCACCTTCTCGCTGCCATC | This paper |
| *nad5*–*nad4* | mitochondrial | F: TCCTTCCACACGAGAGAGTAGG  R: TAACGAACACGGAGACCTTAGC | This paper |
| *rbcL* | chloroplast | F: CTTACCAGYCTTGATCGTTACAAAGG or GCAGCATTTCGTATGACTCC  R: GTAAAATCAAGTCCACCRCG or CTTTCAACTTGGATACCATGAGG | Erickson et al. (2017)^a^ |
| *trnL*–*trnF* | chloroplast | F: CGAAATCGGTAGACGCTACG  R: ATTTGAACTGGTGACACGAG or GGGGATAGAGGGACTTGAAC | Taberlet et al. (1991)^b^; Roslim (2019)^c^ |

^a^Erickson DL, Reed E, Ramachandran P, Bourg NA, McShea WJ, Ottesen A. 2017. Reconstructing a herbivore’s diet using a novel rbcL DNA mini-barcode for plants. AoB Plants 9, plx015.

^b^Taberlet P, Gielly L, Pautou G, Bouvet J. 1991. Universal primers for amplification of three non‐coding regions of chloroplast DNA. Plant Molecular Biology, 17, 1105–1109.

^c^Roslim DI. 2019. Analysis of matK, rbcL and trnL-trnF intergenic spacer sequences on durik-durik (*Syzygium* sp.). Journal of Physics: Conference Series 1351, 012023.

**Supplementary Table 3**. The 118 samples of *Lepidozia* and *Neolepidozia* (outgroup) analysed in this study, with GenBank accession numbers given for the four genetic markers.

| **Species** | **Collection Number** | **Locality** | ***rbcL*** | ***trnL-trnF*** | | | **5.8S rDNA-*ITS2*** | | | ***nad5-nad4*** |
| --- | --- | --- | --- | --- | --- | --- | --- | --- | --- | --- |
| *Lepidozia trichodes* | E.A. Brown 04/67g | Papua New Guinea | JF316324.1 | JF316624.1 | | |  | | |  |
|  | E.A. Brown s.n. | Borneo |  | JX289528.1 | | |  | | |  |
| *Lepidozia laevifolia* | E.A. Brown 04/08 | Australia-NSW | EF101000.1 | EF101109.1 | | |  | | |  |
|  | N.J.M. Gremmen 2000-0385, F | Gough Island |  | JF316622.1 | | |  | | |  |
|  | E.A. Brown 08/260b | New Zealand | JX289389.1 | JX289477.1 | | | JX289220.1 | | |  |
|  | E.D. Cooper 313 | New Zealand |  | JX289520.1 | | | JX289222.1 | | | JX289281.1 |
|  | E.D. Cooper 332 | New Zealand | JX289388.1 | JX289521.1 | | | JX289223.1 | | | JX289283.1 |
|  | E.D. Cooper 264a | New Zealand | JX289390.1 | JX289511.1 | | | JX289221.1 | | | JX289280.1 |
| *Lepidozia bisbifida* | E.D. Cooper 430 | New Zealand | JF316305.1 | JF316627.1 | | |  | | | JX289305.1 |
|  | E.D. Cooper 322 | New Zealand | JX289405.1 | JX289449.1 | | | JX289205.1 | | | JX289307.1 |
|  | E.D. Cooper 311 | New Zealand | JX289406.1 | JX289518.1 | | | JX289206.1 | | | JX289302.1 |
| *Lepidozia ulothrix* | E.A. Brown 08/273 | New Zealand | JF316306.1 | JF316626.1 | | | JX289195.1 | | | JX289317.1 |
|  | E.D. Cooper 116A | Australia-TAS | JX289382.1 | JX289475.1 | | | JX289195.1 | | | JX289317.1 |
|  | E.A. Brown 2012/12 | Australia-TAS | PV730339 | PV730349 | | | PV685808.1 | | | PV730364 |
| *Lepidozia reptans* | N/A (from live culture) | N/A | JF316327.1 | AY608133.1 | | |  | | |  |
|  | (No information) | Finland | GU373419.1 |  | | |  |  |  |  |
|  | Hentschel 01601 | Germany | AM392313.1 |  | | |  | | |  |
| *Lepidozia novae-zelandiae* | E.A. Brown s.n. | New Zealand | JX289392.1 | JX289522.1 | | | JX289218.1 | | | JX289277.1 |
|  | E.A. Brown 08/231 | New Zealand |  | JX289525.1 | | |  | | | JX289285.1 |
|  | E.A. Brown 08/211 | New Zealand | JX289394.1 | JX289478.1 | | | JX289224.1 | | | JX289275.1 |
|  | E.A. Cooper 310 | New Zealand | JX289395.1 | JX289479.1 | | | JX289227.1 | | | JX289284.1 |
|  | E.A. Brown 08/217 | New Zealand | JX289393.1 | JX289524.1 | | | JX289226.1 | | | JX289282.1 |
|  | E.D. Cooper 271 | New Zealand |  | JX289523.1 | | | JX289225.1 | | |  |
| *Lepidozia elobata* | E.A. Brown s.n. | New Zealand | JX289391.1 | JX289527.1 | | | JX289219.1 | | | JX289279.1 |
| *Lepidozia pumila* | E.A. Brown s.n. | New Zealand | JX289387.1 | JX289526.1 | | |  | | | JX289278.1 |
| *Lepidozia concinna* | E.D. Cooper 284 | New Zealand | JX289407.1 | JX289492.1 | | | JX289202.1 | | | JX289304.1 |
|  | E.D. Cooper 424 | New Zealand | JX289412.1 | JX289481.1 | | |  | | | JX289303.1 |
|  | M.A.M. Renner 1077 | New Zealand |  | JX289519.1 | | | JX289201.1 | | |  |
| *Lepidozia bidens* | E.A. Brown 08/200 | New Zealand | JX289377.1 | JX289517.1 | | | JX289216.1 | | | JX289274.1 |
| *Lepidozia eenii* | E.A. Brown s.n. | Australia-NSW | JX289400.1 | JX289516.1 | | | JX289214.1 | | | JX289294.1 |
| *Lepidozia obtusiloba* | E.D. Cooper 309 | New Zealand | JX289397.1 | JX289514.1 | | | JX289184.1 | | | JX289308.1 |
|  | E.D. Cooper 333 | New Zealand | JX289396.1 | JX289515.1 | | | JX289186.1 | | | JX289309.1 |
|  | E.D. Cooper 110 | Australia-TAS | JX289398.1 | JX289513.1 | | | JX289187.1 | | | JX289310.1 |
| *Lepidozia ornata* | E.D. Cooper 296 | New Zealand |  | JX289512.1 | | | JX289185.1 | | | JX289311.1 |
| *Lepidozia acantha* | M.A.M. Renner 4253 | New Zealand | JX289373.1 | JX289510.1 | | | JX289215.1 | | | JX289273.1 |
| *Lepidozia microphylla* | E.D. Cooper 375 | New Zealand | JX289372.1 | JX289458.1 | | | JX289181.1 | | | JX289272.1 |
| *Lepidozia nanophylla* | E.D. Cooper 207 | New Zealand |  | JX289509.1 | | |  | | | JX289289.1 |
| *Lepidozia fugax* | E.D. Cooper 350 | New Zealand | JX289386.1 | JX289480.1 | | | JX289217.1 | | | JX289276.1 |
| *Lepidozia serrulata* | E.D. Cooper 174 | Australia-TAS | JX289384.1 | JX289476.1 | | | JX289197.1 | | | JX289314.1 |
|  | E.A. Brown 08/261 | New Zealand | JX289383.1 | JX289466.1 | | | JX289196.1 | | | JX289316.1 |
|  | J.J. Engel et al. s.n. | New Zealand | PV730342 |  |  |  | PV685799.1 | | | PV730362 |
| *Lepidozia kirkii* | E.D. Cooper 377 | New Zealand | JX289379.1 | JX289474.1 | | | JX289191.1 | | | JX289313.1 |
|  | E.D. Cooper 401 | New Zealand | JX289380.1 | JX289473.1 | | | JX289190.1 | | | JX289269.1 |
|  | E.A. Brown 08/242 | New Zealand | JX289381.1 | JX289467.1 | | | JX289192.1 | | |  |
| *Lepidozia septemfida* | E.D. Cooper 76 | Australia-VIC | JX289375.1 | JX289472.1 | | | JX289182.1 | | | JX289291.1 |
|  | E.D. Cooper 16 | Australia-NSW | JX289376.1 | JX289471.1 | | | JX289183.1 | | | JX289292.1 |
|  | E.A. Brown 2002/118 | Australia-NSW | EF101003.1 | EF101112.2 | | |  | | |  |
|  | M.A.M. Renner 823 | Australia-NSW | EF101004.1 | EF101113.1 | | |  | | |  |
|  | A.L. Rayos 2306 | Australia-VIC | PV730341 | PV730351 | | | PV685810.1 | | | PV730366 |
| *Lepidozia* aff. *septemfida* | M.A.M. Renner 6423 | Australia-QLD | PV730343 | PV730352 | | | PV685802.1 | | | PV730355 |
|  | E.A. Brown 95/218 | Australia-QLD | PV730338 | PV730353 | | | PV685803.1 | | | PV730363 |
|  | D.C. Cargill 73 | Australia-QLD |  |  |  |  | PV685804.1 | | | PV730356 |
|  | J.A. Curnow 4971 | Australia-QLD | PV730344 |  |  |  |  | | | PV730357 |
|  | M.A.M. Renner 7306 | Australia-QLD | PV730345 |  |  |  | PV685805.1 | | | PV730358 |
|  | M.A.M. Renner 7305 | Australia-QLD |  | PV730354 | | | PV685806.1 | | |  |
|  | M.A.M. Renner 6438 | Australia-QLD | PV730346 |  |  |  | PV685807.1 | | | PV730359 |
| *Lepidozia cuspidata* | J.-P. Frahm 15-4 | Chile | JX289374.1 | JX289468.1 | | |  | | |  |
| *Lepidozia hirta* | E.D. Cooper 255 | New Zealand | JX289378.1 | JX289465.1 | | | JX289193.1 | | | JX289315.1 |
| *Lepidozia* aff. *hirta* | T. Thekathyil 217 | Australia-TAS | PV730336 | PV730347 | | | PV685799.1 | | | PV730360 |
|  | T. Thekathyil 206 | Australia-TAS | PV730337 | PV730348 | | | PV685800.1 | | | PV730361 |
|  | A.L. Rayos 2302 | Australia-VIC | PV730340 | PV730350 | | | PV685809.1 | | | PV730365 |
| *Lepidozia pendulina* | E.D. Cooper 421 | New Zealand | JX289416.1 | JX289463.1 | | |  | | | JX289306.1 |
| *Lepidozia spinosissima* | M.A.M. Renner 810 | New Zealand | EF101002.1 | EF101111.1 | | |  | | |  |
|  | E.D. Cooper 243 | New Zealand | JX289371.1 | JX289457.1 | | | JX289180.1 | | | JX289288.1 |
| *Lepidozia glaucophylla* | E.D. Cooper 72 | Australia-VIC | JX289401.1 | JX289438.1 | | | JX289209.1 | | | JX289287.1 |
|  | E.D. Cooper 87 | Australia-VIC | JX289402.1 | JX289437.1 | | | JX289208.1 | | |  |
|  | E.D. Cooper 396 | New Zealand | JX289403.1 | JX289432.1 | | |  | | | JX289286.1 |
| *Lepidozia digitata* | M.A.M. Renner 2547 | New Zealand | JX289411.1 | JX289436.1 | | | JX289213.1 | | | JX289295.1 |
|  | E.D. Cooper 220 | New Zealand | JX289410.1 | JX289434.1 | | |  | | | JX289298.1 |
| *Lepidozia glaucescens* | E.D. Cooper 239 | New Zealand | JX289408.1 | JX289435.1 | | | JX289212.1 | | | JX289296.1 |
|  | E.A. Brown 08/229 | New Zealand | JX289409.1 | JX289433.1 | | | JX289211.1 | | | JX289297.1 |
| *Lepidozia setigera* | J.J. Engel 41 Bog 20, F | New Zealand | JF316326.1 | JF316625.1 | | |  | | |  |
|  | E.D. Cooper 386 | New Zealand |  | JX289426.1 | | | JX289198.1 | | | JX289271.1 |
|  | E.D. Cooper 209 | New Zealand | JX289399.1 | JX289425.1 | | | JX289199.1 | | | JX289270.1 |
| *Lepidozia cladorhiza* | A. Schäfer-Verwimp 24878 | Indonesia | JX289385.1 | JX289423.1 | | |  | | |  |
| *Lepidozia procera* | E.D. Cooper 357 | New Zealand | JX289413.1 | JX289418.1 | | | JX289207.1 | | | JX289300.1 |
|  | E.D. Cooper 381 | New Zealand |  | JX289420.1 | | |  | | | JX289301.1 |
|  | E.A. Brown s.n. | Australia-TAS | JX289414.1 | JX289419.1 | | |  | | | JX289299.1 |
| *Lepidozia cupressina* | D.G. Long 29184 | U.K. |  | JF316623.1 | | |  | | |  |
| *Lepidozia fauriana* | T. Yamaguchi 29076 | Japan |  | JF316620.1 | | |  | | |  |
| *Lepidozia succida* | T. Pocs 97108/D, F | Uganda | KF852317.1 | JF316617.1 | | |  | | |  |
| *Lepidozia vitrea* | H. Miyauchi 269213 | Japan | AB751593.1 |  | | |  | | |  |
| *Lepidozia bragginsiana* | E.A. Brown 08/266 | New Zealand | JX289415.1 | JX289482.1 | | | JX289200.1 | | | JX289293.1 |
| *Lepidozia* undetermined 1a | M. von Konrat, 6/20-4, F | Fiji | JF316325.1 |  | | |  | | |  |
| *Lepidozia* undetermined 2a | B. Shaw 5789, DUKE | China | JF316307.1 |  | | |  | | |  |
| *Lepidozia* undetermined 1b | E.A. Brown s.n. | Borneo |  | JX289491.1 | | | JX289203.1 | | |  |
| *Lepidozia* undetermined 2b | E.A. Brown s.n. | Borneo |  | JX289422.1 | | | JX289204.1 | | |  |
| *Lepidozia* undetermined 3b | E.A. Brown s.n. | Borneo |  | JX289421.1 | | |  | | |  |
| *Lepidozia* undetermined 4b | B. Shaw 8595, DUKE | South Africa |  | JX289470.1 | | | JX289188.1 | | |  |
| *Lepidozia* undetermined 5b | E.D. Cooper 10/001 | Chile |  | JX289469.1 | | | JX289189.1 | | |  |
| *Lepidozia* sp. nov. | E.D. Cooper 269 | New Zealand |  | JX289431.1 | | | JX289210.1 | | |  |
| *Neolepidozia patentissima* | E.A. Brown 03/61 | Australia-VIC | EF101016.1 | EF101125.1 | | |  | | |  |
|  | E.D. Cooper 288 | New Zealand | JX289361.1 | JX289448.1 | | | JX289175.1 | | | JX289262.1 |
|  | E.D. Cooper 162 | Australia-VIC | JX289362.1 | JX289503.1 | | | JX289176.1 | | | JX289253.1 |
| *Neolepidozia heterotexta* | E.A. Brown 03/129 | New Caledonia | EF101012.1 | EF101121.1 | | |  | | |  |
| *Neolepidozia wallichiana* | E.A. Brown 04/829 | Papua New Guinea | JF316328.1 | JF316609.1 | | |  | | |  |
|  | E.A. Brown s.n. | New Zealand | JX289347.1 | JX289508.1 | | | JX289166.1 | | | JX289245.1 |
| *Neolepidozia tetrapila* | J.J. Engel 24292, F | New Zealand | JF316329.1 | JF316611.1 | | |  | | |  |
|  | M.A.M. Renner 2503 | New Zealand | JX289366.1 | JX289502.1 | | | JX289178.1 | | | JX289260.1 |
|  | M.A.M. Renner 2522 | New Zealand | JX289364.1 | JX289455.1 | | |  | | | JX289254.1 |
|  | E.A. Brown 08/215 | New Zealand | JX289365.1 | JX289451.1 | | | JX289179.1 | | | JX289259.1 |
| *Neolepidozia palmata* | E.D. Cooper 107 | Australia-TAS | JX289351.1 | JX289506.1 | | | JX289169.1 | | | JX289255.1 |
|  | E.D. Cooper 146 | Australia-TAS | JX289348.1 | JX289504.1 | | | JX289168.1 | | | JX289257.1 |
|  | E.D. Cooper 149 | Australia-TAS | JX289349.1 | JX289507.1 | | | JX289167.1 | | | JX289256.1 |
|  | E.D. Cooper 169 | Australia-TAS | JX289350.1 | JX289505.1 | | | JX289170.1 | | | JX289250.1 |
| *Neolepidozia seriatitexta* | J.J. Engel 25953, F | Chile | JF316315.1 | JF316621.1 | | |  | | |  |
| *Neolepidozia meridiana* | J.J. Engel 24863, F | New Zealand |  | JF316610.1 | | |  | | |  |
|  | E.D. Cooper 365 | New Zealand | JX289359.1 | JX289452.1 | | |  | | | JX289248.1 |
|  | E.D. Cooper 390 | New Zealand | JX289358.1 | JX289453.1 | | | JX289174.1 | | | JX289258.1 |
| *Neolepidozia tridactylis* | E.D. Cooper 176 | Australia-TAS | JX289352.1 | JX289498.1 | | |  | | | JX289249.1 |
|  | E.D. Cooper 170 | Australia-TAS | JX289355.1 | JX289501.1 | | |  | | |  |
|  | E.D. Cooper 115 | Australia-TAS | JX289353.1 | JX289500.1 | | |  | | | JX289252.1 |
|  | E.D. Cooper 155 | Australia-TAS | JX289354.1 | JX289499.1 | | |  | | |  |
|  | E.D. Cooper 26 | Australia-NSW |  | JX289496.1 | | |  | | |  |
| *Neolepidozia praenitens* | E.A. Brown 08/246 | New Zealand | JX289357.1 | JX289450.1 | | | JX289173.1 | | | JX289251.1 |
|  | E.D. Cooper 336 | New Zealand |  | JX289497.1 | | |  | | |  |
| *Neolepidozia paludicola* | E.D. Cooper 286 | New Zealand | JX289363.1 | JX289456.1 | | | JX289177.1 | | | JX289261.1 |
| *Neolepidozia gibbsiana* | E.D. Cooper 205 | New Zealand | JX289360.1 | JX289454.1 | | | JX289172.1 | | | JX289247.1 |
